# Epigenome-wide meta-analysis of BMI in nine cohorts: examining the utility of epigenetic BMI in predicting metabolic health

**DOI:** 10.1101/2022.07.26.498234

**Authors:** Whitney L. Do, Dianjianyi Sun, Karlijn Meeks, Pierre-Antoine Dugue, Ellen Demerath, Weihua Guan, Shengxu Li, Wei Chen, Roger Milne, Abedowale Adeyemo, Charles Agyemang, Rami Nassir, JoAnn Manson, Aladdin H Shadyab, Lifang Hou, Steve Horvath, Themistocles L. Assimes, Parveen Bhatti, Kristina Jordahl, Andrea Baccarelli, Alicia Smith, Lisa R. Staimez, Aryeh Stein, Eric A. Whitsel, K.M. Venkat Narayan, Karen Conneely

## Abstract

This study sought to examine the association between DNA methylation and body mass index (BMI) and the potential utility of these cytosine-phosphate-guanine (CpG) sites in predicting metabolic health. We pooled summary statistics from six trans-ethnic EWAS of BMI representing nine cohorts (n=17058), replicated these findings in the Women’s Health Initiative (WHI, n=4822) and developed an epigenetic prediction score of BMI. In the pooled EWAS, 1265 CpG sites were associated with BMI (p<1E-7), and 1238 replicated in the WHI (FDR < 0.05). We performed several stratified analyses to examine whether these associations differed between individuals of European descent and individuals of African descent. We found five CpG sites had a significant interaction with BMI by race/ethnicity. To examine the utility of the significant CpG sites in predicting BMI, we used elastic net regression to predict log normalized BMI in the WHI (80% training/20% testing). This model found 397 sites could explain 32% of the variance in BMI in the WHI test set. Individuals whose methylome-predicted BMI overestimated their BMI (high epigenetic BMI) had significantly higher glucose and triglycerides, and lower HDL-cholesterol and LDL-cholesterol compared to accurately predicted BMI. Individuals whose methylome-predicted BMI underestimated their BMI (low epigenetic BMI) had significantly higher HDL-cholesterol and lower glucose and triglycerides. This study identified 553 previously identified and 685 novel CpG sites associated with BMI. Participants with high epigenetic BMI had poorer metabolic health suggesting that the overestimation may be driven in part by cardiometabolic derangements characteristic of metabolic syndrome.

## Introduction

Globally, the prevalence of obesity is rising with an estimated 650 million adults obese, representing 19.5% of the adult population [1, 2]. Obesity has been found to accompany a multitude of molecular and metabolic perturbations including impaired cell signaling, insulin resistance, hyperlipidemia, and hypertension [3-5]. Ultimately these perturbations can lead to the early onset of chronic diseases with individual living with obesity having a 37% increased risk of type 2 diabetes [6] and 67-85% increased risk of cardiovascular disease compared to individuals living without obesity [7]. With a growing population of individuals living with obesity, it is increasingly important to understand the molecular mechanisms dysregulated by obesity to further elucidate both early markers of disease progression and novel therapeutic targets.

Epigenetic mechanisms are molecularly-mediated changes in gene function which do not change the DNA sequence. DNA methylation, the most widely characterized epigenetic mechanism, occurs when a methyl group attaches to the cytosine in a cytosine-guanine nucleotide (CpG) pair [8]. DNA methylation has been shown to influence gene expression by blocking transcription factor binding and recruiting chromatin remodelers [9]. As a functional mechanism influencing gene expression, DNA methylation may be on a disease pathway and could provide insight into important therapeutic targets. DNA methylation has also become an important biomarker of health, for example with the development of epigenetic clocks, which can provide accurate estimates of individual age based on the methylation status of a representative set of CpG sites [10]. Individuals whose DNA methylation deviate from their actual chronological age, such that their epigenetically predicted age is higher than their actual age, have been shown to have higher rates of cancer, cardiovascular disease, diabetes, and mortality [11]. All of these properties may be relevant in the relationship between DNA methylation and obesity.

Several studies have examined the relationship between DNA methylation and body mass index (BMI), a commonly used measure of obesity [12-21]. Obesity has been significantly associated with differential DNA methylation, and Mendelian randomization analyses have suggested that while this differential methylation appears to be a consequence of the state of obesity at many CpG sites, some CpGs show evidence consistent with causal roles in obesity [13, 18]. While several large-scale studies have identified sites associated with obesity, it is likely that additional sites will be detectable only with large sample sizes, as observed DNA methylation differences are often subtle [22]. Thus, a goal of this study is to conduct the largest epigenome-wide association study (EWAS) meta-analysis of BMI in nine population-based cohort studies to identify novel sites associated with obesity. The identification of novel sites can reveal unique molecular signatures of various BMI phenotypes (including metabolically healthy/unhealthy BMI) and may enable improved prediction of BMI. Previous studies have reported that a collection of methylation-based predictors can explain between 4.7-18% of the variance in BMI [13, 21, 23, 24]. In conducting the largest EWAS, we may have better predictive capacity by incorporating the novel CpG sites identified in the EWAS meta-analysis. As such, a secondary aim of this study is to examine whether BMI-associated CpG sites can predict BMI. As with epigenetic age, deviations from epigenetically predicted BMI may be associated with several relevant health outcomes and could be used as an informative metric of overall health and/or a predictor of future cardiovascular disease. Thus, we examined whether individuals whose BMI was poorly predicted by DNA methylation (DNA methylation over predicts their actual BMI or DNA methylation under predicts their actual BMI) have differential metabolic health status.

## Methods

### Participants

Our discovery analysis used data from 17,034 participants from six published EWAS studies of individuals of European descent (n=11220), African descent (n=2587), and South Asian descent (n=2680). The six studies were based on nine cohorts: Atherosclerosis Risk in Communities study (ARIC) [25], Melbourne Collaborative Cohort Study (MCCS) [26], Lifelines DEEP [27], Lothian Birth Cohort (LBC) 1921 and 1936 [28], Bogalusa Heart Study (BHS) [29], the Research on Obesity and Diabetes among African Migrants (RODAM) [30], the Kooperative Gesundheitsforschung in der Region Augsburg (KORA) [31], the London Life Sciences Prospective Population Study (LOLIPOP) [32], and Italian cardiovascular component of the European Prospective Investigation into Cancer and Nutrition (EPICORE) [33]. Replication analyses were conducted in three ancillary studies from the Women’s Health Initiative (WHI): *Epigenetic Mechanisms of Particulate Matter-Mediated Cardiovascular Disease* (EMPC, aka AS315), the *Integrative Genomics for Risk of Coronary Heart Disease and Related Phenotypes in WHI cohort* (BAA23), and *Bladder Cancer and Leukocyte Methylation* (AS311). In the WHI, individuals were excluded if BMI and blood samples for DNA methylation were not measured within the same year. Extreme levels of BMI <17 kg/m^2^ and >75 kg/m^2^ were excluded. Further description of the discovery and replication cohorts is described in **Supplemental Methods**.

### BMI, DNA methylation and covariates

BMI was defined as weight in kg/height in m^2^. Methodologies obtaining weight and height differed among the studies, however all used standard methods. One study transformed BMI values to obtain a normal distribution [20]. Relevant variables in our replication analysis included race/ethnicity, age, physical activity and smoking status. Race/ethnicity, smoking and physical activity were self-reported. Smoking status was defined as current, former or never.

DNA methylation was measured in several cell types including CD4+ T-cells, mononuclear cells and whole blood. DNA methylation in all studies was measured using the Illumina 450K Infinium Methylation BeadChip. DNA methylation was estimated as the proportion of methylated signal relative to combined unmethylated and methylated signal for a specific CpG site, defined as the β-value. Quality control procedures of the previous studies have been reported in detail and they did not differ substantially across studies. In the WHI, all methylation data were quality controlled and normalized using beta-mixture quantile normalization. In replication analyses, chip and row were included as technical covariates in all models to adjust for batch effects. Cell composition was estimated using methods derived by Houseman et al. [34].

### Statistical Analysis

A summary of our analyses is included in **Figure 1A**. Our primary method was weighted sum of Z-score meta-analysis [35]. This method utilizes Z-scores from individual study summary statistics computed from inverse-normal p-values and the direction of effect to determine significant sites. This was chosen as the primary method for meta-analysis since the studies did not all have equivalent exposure-outcome definition (DNA methylation defined as exposure in two studies and outcome in four studies) and BMI was transformed in one study. The EWAS was adjusted for genomic inflation and significance was defined as p < 1×10^−7^.

**Figure 1.**
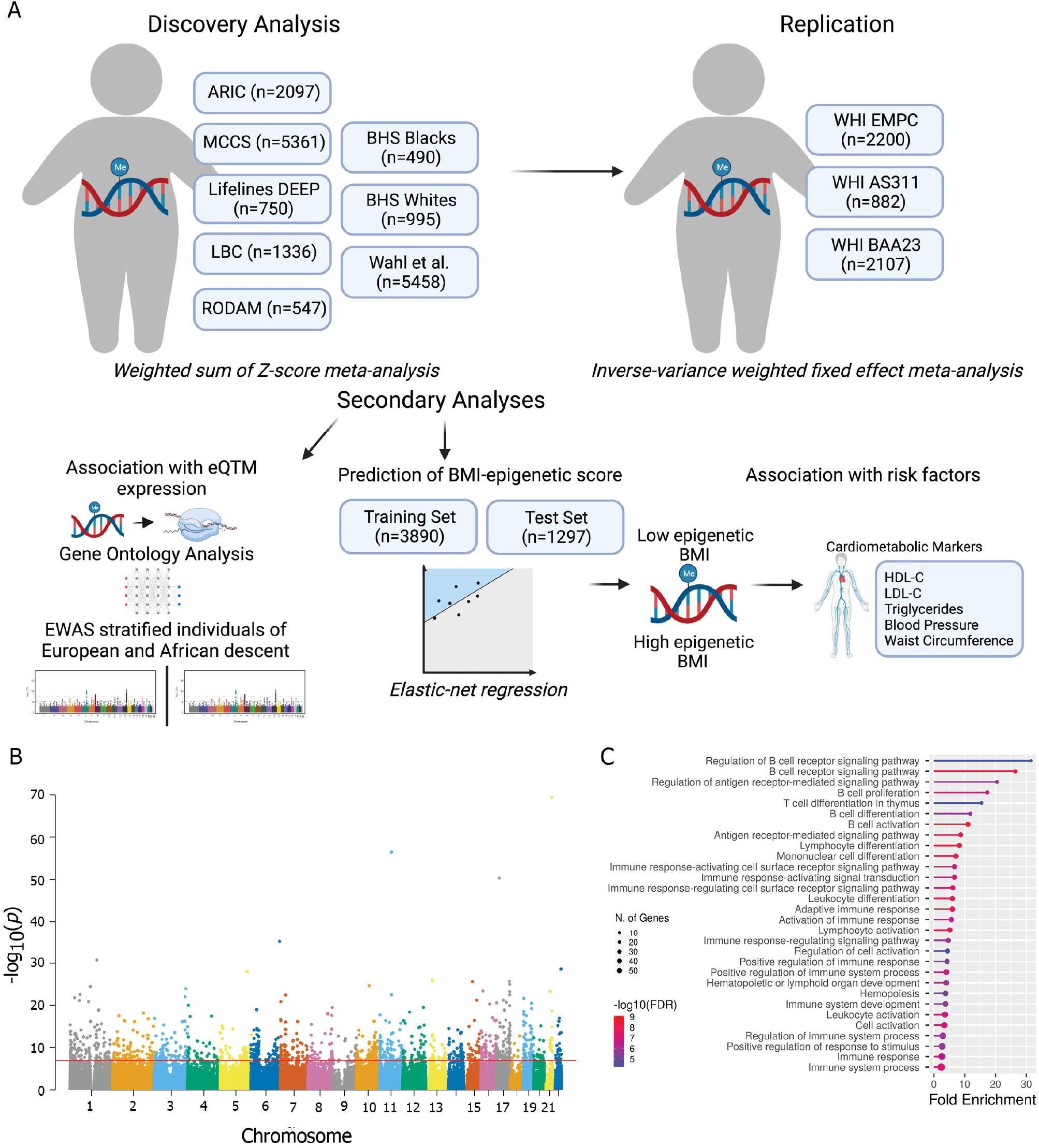
1A Description of study analyses. 1B Manhattan plot of the association between DNA methylation and BMI. 1C Top pathways identified in gene ontology analysis.

The significant sites were examined for replication within WHI. Models were stratified by ancillary study. Covariates in this analysis included age, race/ethnicity, cell composition, the top three principal components of genetic relatedness, smoking status, clinical trial arm and case-control status (BAA23 and AS311). To account for potential chip-to-chip differences in measurement and to adjust for batch effects, chip was included as a random effect for each BeadChip in our model. Stratified analyses were combined using inverse-variance weighted (IVW) meta-analysis [36]. Significance was defined by false discovery rate (FDR) q-value < 0.05.

### BMI Prediction Score

To examine the degree to which methylation can predict BMI and the secondary cardiometabolic outcomes associated with BMI, we used elastic net regression models with the significant sites to predict log-normalized BMI. The WHI cohorts were randomly divided into a training and test set (80% and 20%, respectively) with an equal BMI distribution. We used elastic net regression on the training set with 10-fold cross validation to select a predictive model, which we subsequently tested in the test set. Using the significant sites and coefficients selected by the model, a DNA methylation prediction score was developed by multiplying the coefficient by the individual β-value and summing over all the sites for each individual. We then evaluated the performance of the DNA methylation score in the test set, both in terms of how accurately it predicted BMI (metrics: *R*^*2*^ and median absolute deviation) and how well it predicted obesity status (BMI ≥ 30 kg/m^2^) (metrics: sensitivity and specificity).

Using the predicted BMI values, we examined the patterns among outliers in the prediction model. Individuals were split into categories based on regressing the predicted BMI on the actual BMI. Accurately predicted individuals were defined as those with residuals between -0.04 to 0.04 (accurate epigenetic BMI). Individuals outside of this range were split into two groups: residual below -0.04 (low epigenetic BMI or individuals whose methylome-predicted BMI underestimated their BMI) and residual above 0.04 (high epigenetic BMI or individuals whose methylome-predicted BMI overestimated their BMI). These thresholds were defined based on the 10% and the 90% distribution of the residuals. Using these categories, we examined cardiometabolic differences including waist circumference, triglycerides, HDL-cholesterol, LDL-cholesterol, and blood glucose among these categories using linear regression models regressing log-normalized cardiometabolic markers on DNA methylation prediction category adjusted for age, race/ethnicity, smoking status and physical activity. To aid interpretability, results were reported based on the change in average value in the text. We additionally examined results using thresholds defined by the 20% and 80% distribution of residuals and found consistent findings.

### Sensitivity Analyses

We conducted several sensitivity analyses. In the discovery meta-analysis, we examined the influence of specific studies on the results using leave-one out analyses to examine the degree that each study is influencing the results. We also compared results from the weighted sum of z-score meta-analysis with results that would be obtained using an IVW meta-analysis in studies with the same exposure-outcome definition. We examined several sites for interaction by self-reported race/ethnicity and BMI using linear mixed-effect models adjusting for age, cell composition, smoking status, WHI study randomization arm, case-control status, row with a random effect for chip. In our replication analysis in WHI, models were additionally adjusted for diet quality, physical activity level and socioeconomic status.

## Results

Our discovery analysis included 17058 participants from six EWAS (**Figure 1A, Table 1** and **Supplementary Table 1**). The definition of BMI and DNA methylation differed with several transforming these values in the models (**Table 1**). The covariates in the model also differed with all studies adjusting for age and sex, and the majority adjusting for cell composition and smoking status. When pooling results from all studies, 1265 CpG sites were associated with BMI (**Figure 1B, Supplementary Table 2**, p < 1E-7) with 498 of the sites having a consistent direction of effect in all of the cohorts meta-analyzed. More than half of the significant sites (726 CpG sites) were positively associated with BMI.

**Table 1.**
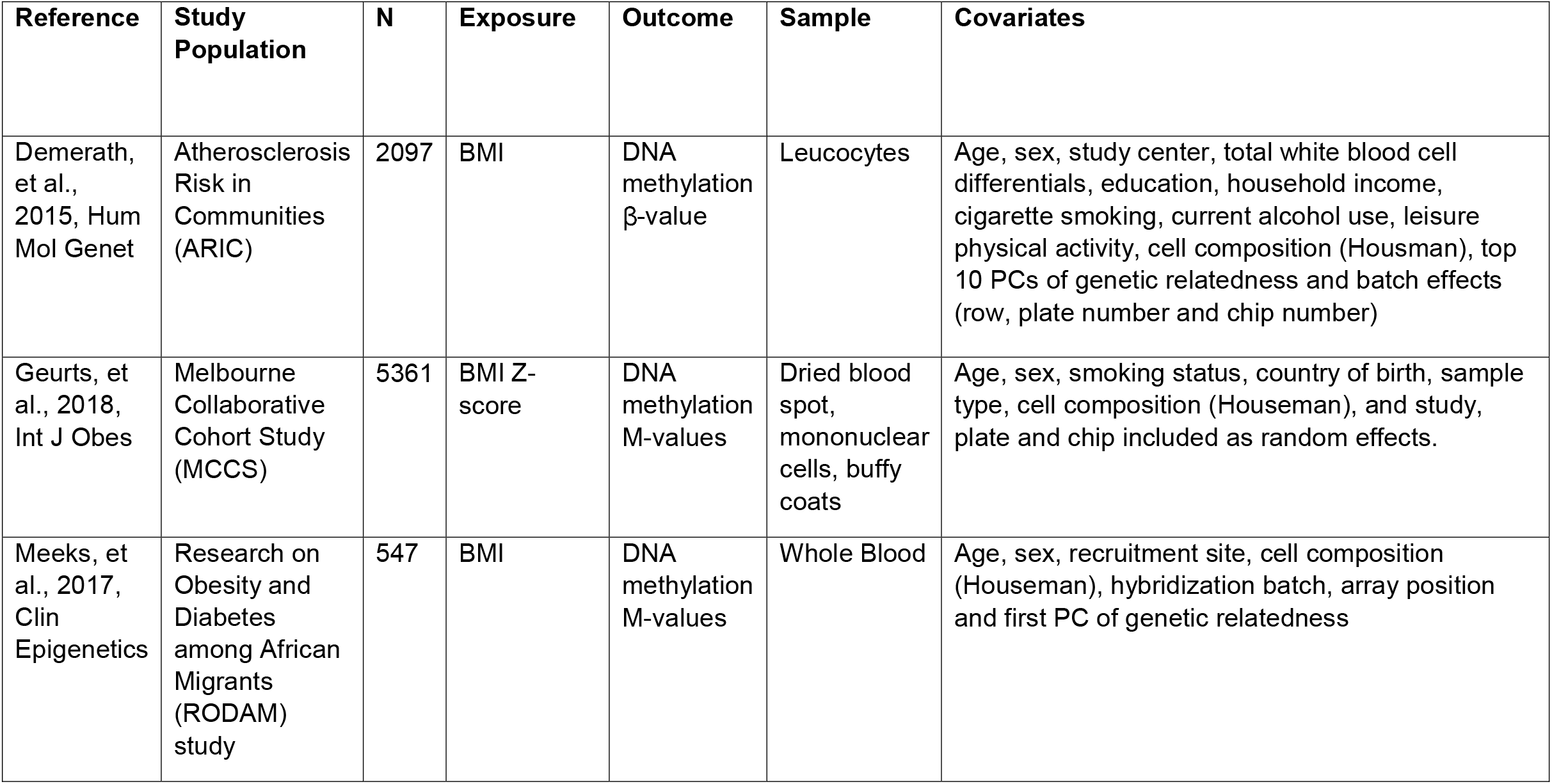
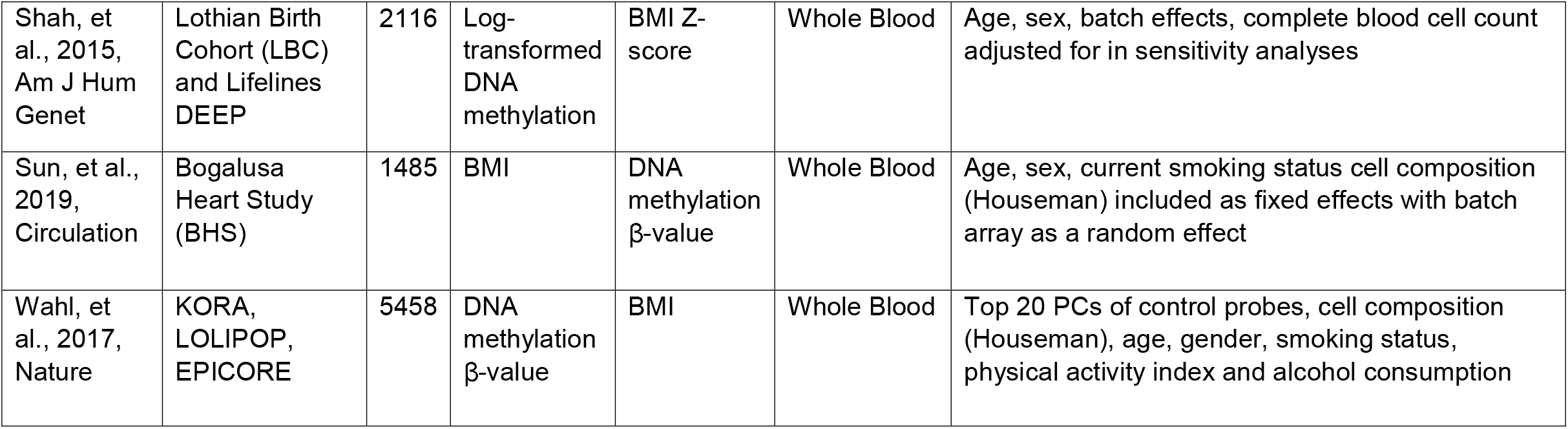
Study characteristics of discovery analyses.

In the WHI, 367 women were excluded due to missing BMI, extreme levels of BMI, or overlap leaving 4822 women included in the replication cohort (**Supplementary Table 3**). Of the 1265 sites identified in the discovery analysis, 1254 were analyzed after QC. In the WHI, 1238 CpG sites were significantly associated with BMI (**Supplementary Table 4**, FDR q-value < 0.05). These 1238 CpG sites annotated to 742 unique genes. Additionally, 147 of these genes were annotated to more than one BMI-associated CpG site, with 382 CpG sites annotated to these 147 genes. With the large sample size, we were able to discover 685 novel CpG sites that had not previously been identified in EWAS of BMI as well as 553 CpG sites previously identified in the literature. We examined how the replicated sites associated with differential gene expression based on previously published analyses of the Grady Trauma Project (GTP) and Multi-Ethnic Study of Atherosclerosis (MESA) cohort [37]. The 1238 CpG sites associated with 1103 CpG-mRNA associations in MESA (**Supplementary Table 5**) and 79 CpG-mRNA associations in GTP (**Supplementary Table 6**). One site associated with the same mRNA transcript in both cohorts, cg25653947, which was positively associated with expression in *TOP1MT*. We performed a gene ontology (GO) analysis of the differentially expressed genes and found enrichment in pathways related to the adaptive immune system with regulation in B- and T-cell pathways (**Figure 1C, Supplementary Table 7**).

We next re-performed our discovery EWAS stratified by European vs. African descent. We found 936 and 130 CpG sites that were associated with BMI in the analyses restricted to individuals from European (n=11,220) and African (n=2,587) descent, respectively. Of the 130 significant CpG sites in the analysis of individuals of African descent, 43 unique sites were only significant in that population (**Supplementary Table 8-9**). We examined these sites for interaction in the WHI non-Hispanic white and African American individuals. We found that five CpG sites had a significant interaction with BMI by race/ethnicity (**Table 2, Supplementary Fig.1**). Two sites were quantitative trait methylation loci in the GTP cohort: cg25212453 negatively associated with *TNFRSF13B* and *COCH* and cg08122652 negatively associated with *LGALS3BP* and *OTOF* (**Supplementary Table 10**).

**Table 2.** Interaction between BMI and race/ethnicity in WHI between non-Hispanic whites and African Americans.

| CpG Site | Effect Estimate | Standard Error | Z-score | P-value |
| --- | --- | --- | --- | --- |
| cg25652701 | -5.8E-04 | 1.4E-04 | -4.29 | 1.76E-05 |
| cg25212453 | 5.01E-04 | 1.6E-04 | 3.05 | 0.002 |
| cg08122652 | 8.9E-04 | 3.6E-04 | 2.47 | 0.014 |
| cg27113059 | -2.3E-04 | 9.59E-05 | -2.36 | 0.018 |
| cg15391590 | -2.4E-04 | 1.1E-04 | -2.18 | 0.029 |

**Table 3.**
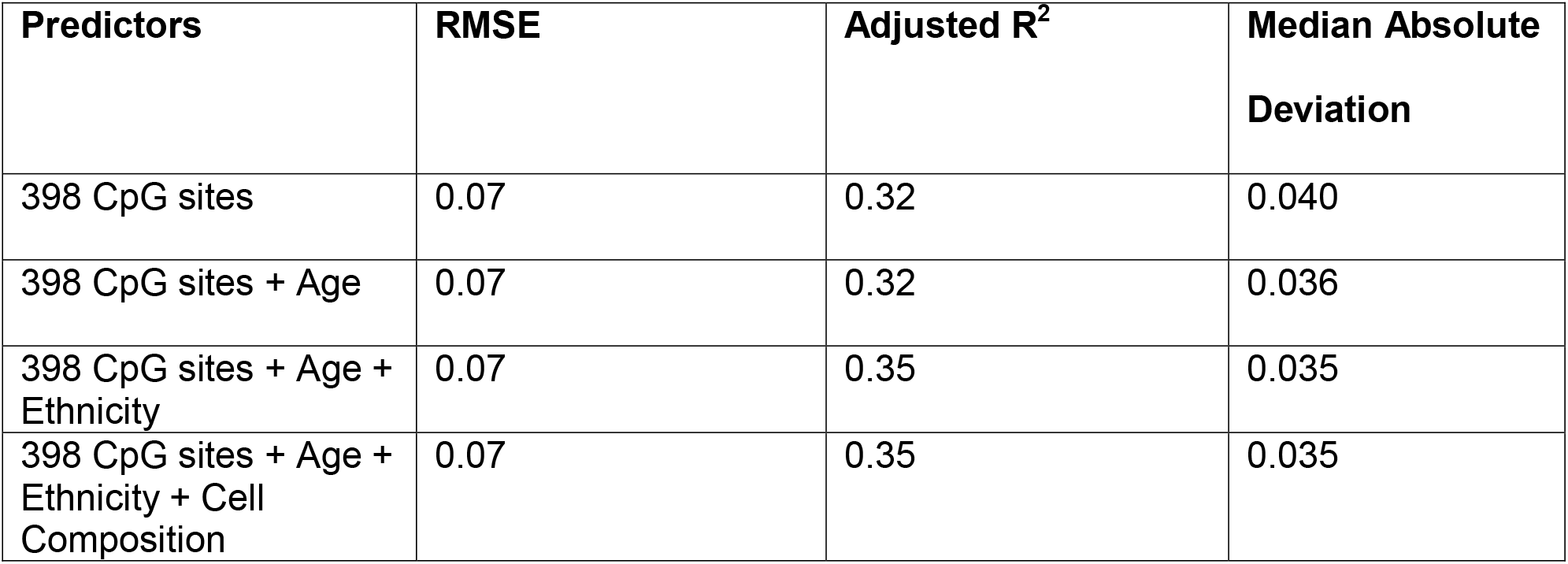
Predicting BMI from DNA methylation using elastic net regression

| Predictors | RMSE | Adjusted R<sup>2</sup> | Median Absolute Deviation |
| --- | --- | --- | --- |
| 398 CpG sites | 0.07 | 0.32 | 0.040 |
| 398 CpG sites + Age | 0.07 | 0.32 | 0.036 |
| 398 CpG sites + Age + Ethnicity | 0.07 | 0.35 | 0.035 |
| 398 CpG sites + Age + Ethnicity + Cell Composition | 0.07 | 0.35 | 0.035 |

**Table 4.**
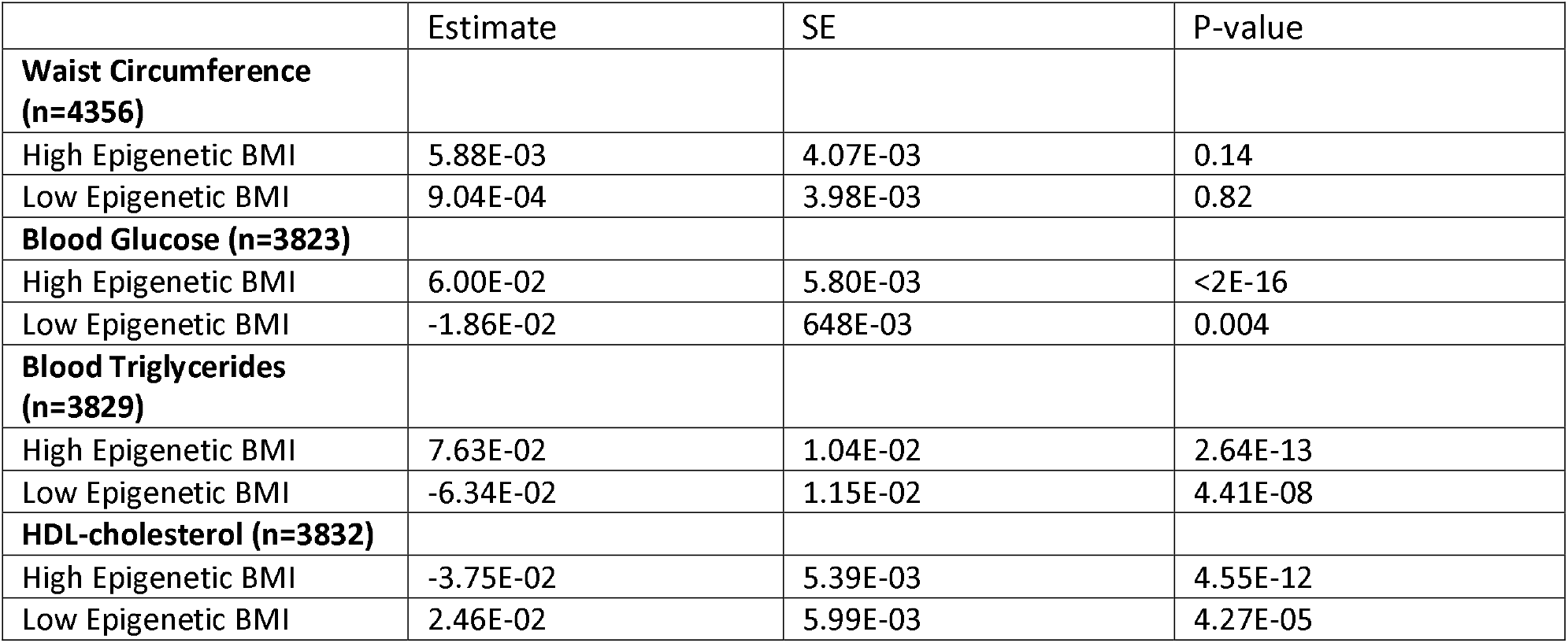
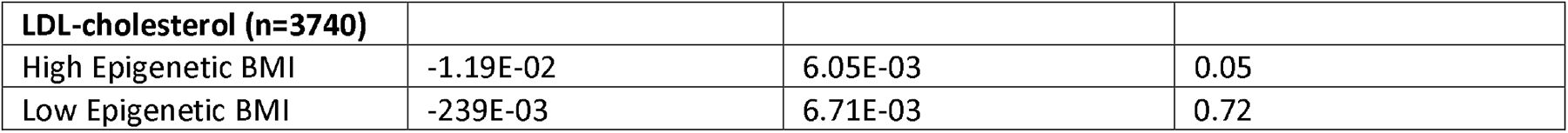
Outliers in the prediction model compared to log-normalized cardiometabolic risk factors. Model adjusted for race/ethnicity, smoking status, age and physical activity.

We next explored the potential of DNA methylation to predict BMI using the 1238 CpG sites from the replication analysis. After model tuning using elastic-net regression in a training set (N=3858), 398 sites were selected for the model (**Supplementary Table 11**). These sites accounted for 32% of the variance in BMI in the test set (MAD = 0.04, N=964). The addition of age, race/ethnicity, physical activity, and cell composition as predictors only marginally improved the adjusted R^2^ (**Table 4**). In the combined training and test set (N=4822), these sites accounted for 36% of the variance in BMI. For comparison we examined how well the predictors from Mendelson *et al*. [13] estimated BMI in the WHI cohort. In the full WHI cohort, the 83 CpG sites accounted for 29% of the variance.

We next assessed the potential of this DNA methylation-based BMI score to predict obesity, defined as BMI>30. In our test set (N=964), the sensitivity was 0.82 and the specificity was 0.57 with an area under the curve of 0.69 (**Figure 2A**). Individuals were then categorized based on how well methylation predicted BMI. On average, DNA methylation tended to underpredict BMI (**Figure 2B**). Individuals with high epigenetic BMI had 20.5 mg/dL higher blood glucose (SE: 2.0, p<2E-16), 31 mg/dL higher triglycerides (SE: 4.3, p=9.24E-08), 4.3 mg/dL lower HDL-cholesterol (SE: 0.68, p=1.06E-07) and 3.3 mg/dL lower LDL-cholesterol (SE: 2.0, p=0.047) compared with accurate predicted BMI. In contrast, individuals with low epigenetic BMI had 5.2 mg/dL lower blood glucose (SE: 2.2, p<2E-16), 23.7 mg/dL lower triglycerides (SE: 4.8, p=2.39E-08) and 3.0 mg/dL higher HDL-cholesterol (**Figure 2C**, SE: 0.8, p=0.0004) compared to accurate predicted BMI.

**Figure 2.**
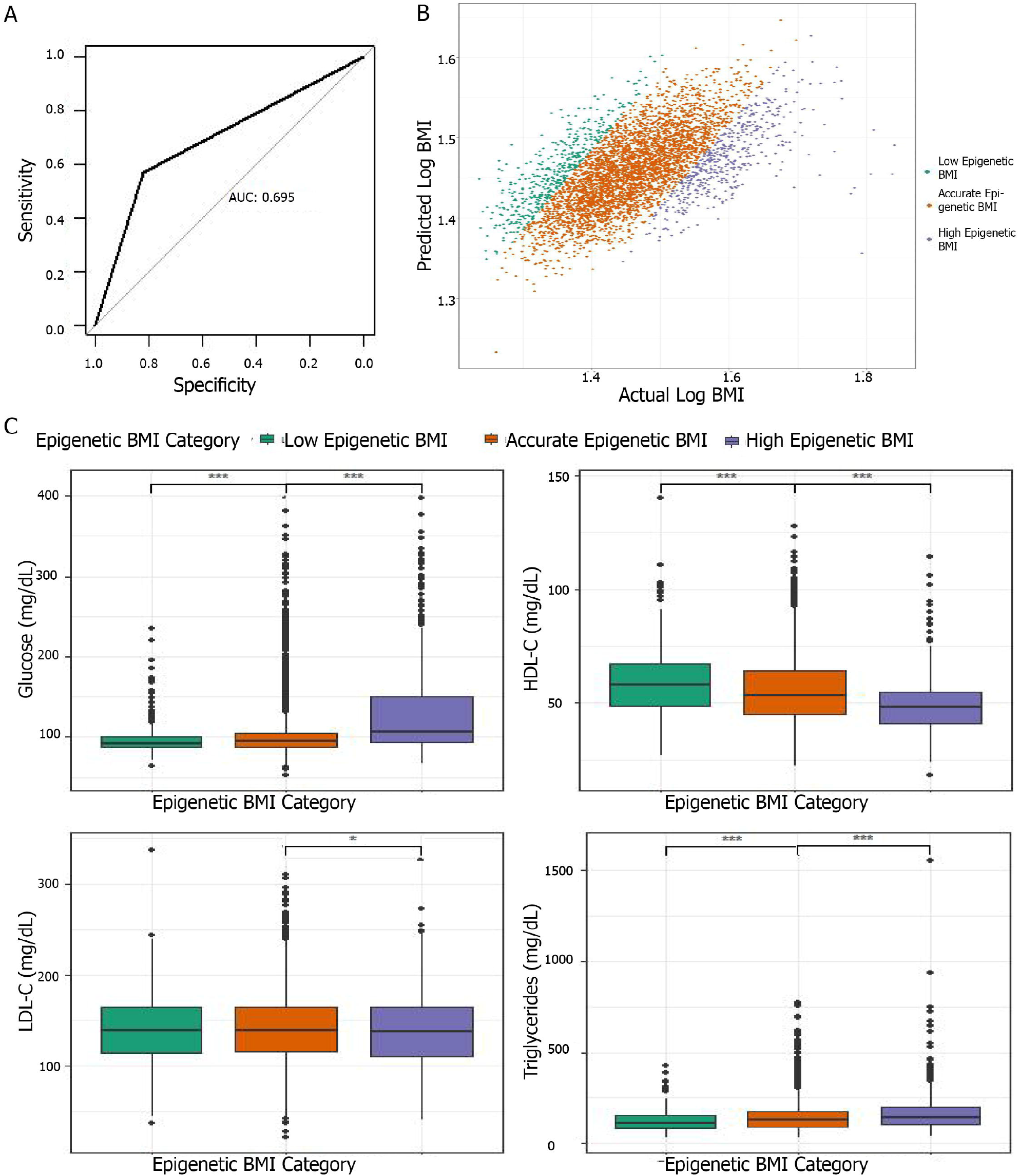
2A Receiver operating characteristic curve showing the performance of the DNA methylation prediction score identifying obesity. AUC denotes area under the curve. Y-axis is the sensitivity (true positive rate) and the x-axis is 1-specificity (false positive rate). 2B Scatter plot of predicted BMI from elastic net regression of 398 CpG sites by actual BMI. Individuals categorized based on the residual of predicted BMI regressed on actual BMI. 2C Boxplot of the association between epigenetic prediction category and blood glucose (mg/dL), high density lipoprotein (HDL-C, mg/dL), low density lipoprotein (LDL-C, mg/dL) and triglycerides (mg/dL).

We conducted several sensitivity analyses. We first examined how the results changed in a leave-one out meta-analysis (**Supplementary Table 12**). Excluding the results from Wahl *et al*. [18] led to the largest reduction in significant sites resulting in 536 significant CpG sites. However, this is likely due to a reduction in power. We next compared results obtained using Z-score vs. IVW meta-analysis in cohorts with the same exposure-outcome relationship (ARIC, RODAM, BHS White and BHS Black). In the IVW and Z-score meta-analysis of these four studies, 1939 CpG sites and 1433 CpG sites, respectively, were significantly associated with BMI (p < 1E-7) with 935 overlapping sites among methods. Among the sites identified significant in either analysis, the correlation between the test statistics obtained using Z-score vs. IVW meta-analysis was 0.98. The meta test statistics tend to be smaller when identified using weighted sum of Z-score meta-analysis, suggesting our main results may be more conservative than what would be obtained using IVW meta-analysis. Finally, in the main EWAS, we examined how the results changed when adjusted for diet, physical activity and income status. Overall, 1161, 1167 and 1160 CpG sites remained associated with BMI when additional covariates were included to adjust for diet quality, physical activity and income, respectively.

## Discussion

This study identified a unique methylomic signature of BMI and obesity. In the WHI, the majority of the sites identified in the discovery cohort (99%) were replicated and found to predict several metabolic and inflammatory pathways. Moreover, we found five CpG sites that are differentially associated with BMI between non-Hispanic whites and African Americans, two of which may play a role in gene expression. Finally, we constructed a score based on 398 CpG sites that was able to predict BMI as well as several other cardiometabolic risk factors. Individuals whose measured BMI was higher than predicted by their methylome were found to have poorer metabolic health including higher blood glucose and triglycerides and lower HDL-cholesterol compared to individuals whose BMI was accurately predicted.

This study identified 1238 CpG sites that were significantly associated with BMI in several race/ethnicity groups. The 1238 CpG sites were associated with differential gene expression in MESA and GTP. In the GO analysis of the differentially expressed transcripts, the most significant enriched pathways were immune response pathways, particularly the adaptive immune response. The top pathways regulated B- and T-cell signaling. Low-grade inflammation in obesity is a hallmark of the disease, which leads to significant metabolic dysregulation [38]. Several studies have found BMI-associated CpG sites are enriched for immune pathways [13, 39]. Mechanistic studies have identified DNA methylation as playing a key role in promoting macrophage polarization in response to obesity, with more M1 macrophages associated with obesity [40].

We were also able to examine how these associations differed when stratified by race/ethnicity. Racial and ethnic differences in adiposity have been well established. While African Americans have been found to have higher risk for cardiovascular diseases compared to non-Hispanic whites, they have consistently been shown to have lower visceral adipose tissue and lower body fat percentage compared to non-Hispanic whites [41, 42]. Of the five sites with a significant interaction between BMI and race/ethnicity, two CpG sites were associated with differential expression in four mRNA transcripts. These gene transcripts are related to inflammatory pathways and hearing. *TNFRSF13B* and *LGALS3BP* were differentially expressed in association with two CpG sites. These two genes have been found to be regulators of NF-kappa-B signaling and to be upregulated with obesity [43-45]. Our study found a positive association between BMI and methylation in cg25212453 and cg08122652 (in WHI) and a negative association between methylation in these two sites and expression in *LGALS3BP* and *TNFRSF13B* (in GTP). Thus, as BMI increases in African Americans, gene expression may be decreasing in these sites, suggesting a potentially advantageous effect on inflammatory profiles in African Americans. Low-grade inflammation in obesity leads to significant metabolic dysregulation [38]. However, there is some epidemiological data that suggests individuals of African descent may not be as prone to an increased inflammatory profiles when living with obesity [46, 47]. Our study may provide some mechanistic explanation to these differences in the relationship between inflammation and adiposity in individuals of African descent.

We also found that DNA methylation was predictive of BMI, with the score we developed based on 398 CpG sites explaining 32% of the variance in BMI in an independent test set. Previous studies constructing scores based on smaller samples have been able to explain between 4.7-18% of the variance in BMI [13, 21, 23, 24]. DNA methylation has been found to be an accurate predictor of current BMI and a poor predictor of future BMI [24]. Outliers in the epigenetic BMI model predicted a unique phenotype. Individuals with high epigenetic BMI or whose BMI was over-predicted by the epigenetic markers had poorer cardiometabolic markers compared to accurately predicted BMI. This may suggest that epigenetic BMI prediction may be identifying individuals with poor health regardless of their BMI and that these sites may be useful biomarkers to examine further. Our findings related to LDL-C were inconsistent with other cardiometabolic markers. We found that individuals with high epigenetic BMI had lower LDL-C as compared to individuals with accurate epigenetic BMI. These results were verging on null (p=0.0497), thus they should be interpreted with caution.

Some of our findings should be interpreted with caution given several important limitations. In the discovery analysis, we stratified this analysis based on race/ethnicity as it was defined within each of the individual studies. This differed between studies with most based on self-report of race/ethnicity. Thus, it is unclear whether we are identifying molecular differences due to ancestry or social construct. Moreover, these populations, which include African Americans, Ghanaians, and European-residing Ghanaians, are not homogenous in genetic ancestry, living environment, lifestyles and other factors. Nevertheless, our interaction and expression analyses were conducted in African American populations from the WHI and GTP, so these results may only be generalizable for this population. In particular, the racial disparities in the US may be an underlying cause of these results, as opposed to differences in ancestry. For example in the US, African Americans are much more likely to live in poverty compared to non-Hispanic whites [48]. In our results, we may be identifying compensatory mechanisms of structural racism which may be driven by environmental exposures for example, ambient particulate matter exposure, stress, lack of access to health care as well as obesity. Another potential limitation is that the training and test set in our prediction analyses come from the same population (WHI). Future research efforts could test this model in another population to examine the reproducibility of these findings.

Overall, this study yields several important discoveries. We identified novel sites associated with BMI and found a unique molecular profile associated with obesity in individuals of African descent. We additionally found that epigenetic markers can predict BMI well and may be able to distinguish individuals whose metabolic health does not align with their BMI. Future studies should examine whether BMI-associated methylation is differential by metabolic health status.

## Acknowledgments

We would like to acknowledge and thank Drs. Sonia Shah and Peter Visscher for sharing summary statistics which have been included in the discovery portion of this analysis. The WHI program is funded by the National Heart, Lung, and Blood Institute, National Institutes of Health, U.S. Department of Health and Human Services through 75N92021D00001, 75N92021D00002, 75N92021D00003, 75N92021D00004, 75N92021D00005. WHI EMPC (AS315) was supported by NIEHS grant R01-ES020836 (LH, AB, EAW). WHI AS311 was supported by American Cancer Society award 125299-RSG-13–100-01-CCE (PB). WHI-BAA23 was supported by NHLBI Broad Agency Announcement contract HHSN268201300006C. The WHI program is funded by the National Heart, Lung, and Blood Institute, National Institutes of Health, U.S. Department of Health and Human Services through contracts 75N92021D00001, 75N92021D00002, 75N92021D00003, 75N92021D00004, 75N92021D00005. We would acknowledge the following WHI investigators: Jacques Rossouw, Shari Ludlam, Joan McGowan, Leslie Ford, and Nancy Geller (National Heart, Lung, and Blood Institute, Bethesda, Maryland); Garnet Anderson, Ross Prentice, Andrea LaCroix, and Charles Kooperberg (Fred Hutchinson Cancer Research Center, Seattle, WA); Barbara V. Howard (MedStar Health Research Institute/Howard University, Washington, DC); Marcia L. Stefanick (Stanford Prevention Research Center, Stanford, CA); Rebecca Jackson (The Ohio State University, Columbus, OH); Cynthia A. Thomson (University of Arizona, Tucson/Phoenix, AZ); Jean Wactawski-Wende (University at Buffalo, Buffalo, NY); Marian Limacher (University of Florida, Gainesville/Jacksonville, FL); Jennifer Robinson (University of Iowa, Iowa City/Davenport, IA); Lewis Kuller (University of Pittsburgh, Pittsburgh, PA); Sally Shumaker (Wake Forest University School of Medicine, Winston-Salem, NC); Robert Brunner (University of Nevada, Reno, NV); and Mark Espeland (Wake Forest University School of Medicine, Winston-Salem, NC). The RODAM study was supported by the Intramural Research Program of the National Human Genome Research Institute of the National Institutes of Health (NIH) through the Center for Research on Genomics and Global Health (CRGGH) and by the European Commission under the Framework Programme (Grant Number: 278901). The CRGGH is also supported by the National Institute of Diabetes and Digestive and Kidney Diseases and the Office of the Director at the NIH (Z01HG200362). The Atherosclerosis Risk in Communities study has been funded in whole or in part with Federal funds from the National Heart, Lung, and Blood Institute (NHLBI), National Institutes of Health, Department of Health and Human Services (contract numbers HHSN268201700001I, HHSN268201700002I, HHSN268201700003I, HHSN268201700004I, and HHSN268201700005I). The authors thank the staff and participants of the ARIC study for their important contributions. Funding was also supported by 5RC2HL102419 and R01NS087541. WD and KMVN were supported in part by P30 Georgia Diabetes Translation Research Center and RADx-UP Diabetes. KMVN was partly supported by the National Institute Of Diabetes And Digestive And Kidney Diseases of the National Institutes of Health under Award Number P30DK111024 and P30111024-05S1. The content is solely the responsibility of the authors and does not necessarily represent the official views of the National Institutes of Health. PAD was supported in part by NHMRC grant 1164455. RM was supported in part by NHMRC grant 1088405. AA was supported in part by NIH IRP - 1ZIAHG200362. TA was supported in part by NHLBI contract HHSN268201300006C. PB was supported in part by American Cancer Society (125299-RSG-13-100-01-CCE). KJ was supported in part by T32-CA094880.

## Author Contributions

WLD, KVN, and KNC conceived of the study. WLD, KVN, LRS, ADS and AS developed the methods. ED, WG, SL, WC, DS, PMV, SS, RM, PAD, KM, AA, and CA computed and shared the summary statistics from the participating studies in the discovery analysis. LH, SH, TA, PB, KMJ, EAW and AB curated and processed the data used in the replication analyses. WLD performed all statistical analyses with support from KNC. WLD drafted the manuscript with content expertise and review from all authors. All authors read and approved of the final manuscript.

## Data and Code Availability

All datasets used in this study are publicly available at individual study websites. The code used to generate these findings are available from the corresponding author upon reasonable request.

