## Supplemental Methods for "Epigenome-wide meta-analysis of BMI in nine cohorts: examining the utility of epigenetic BMI in predicting metabolic health"

**Supplemental Materials**

**Supplemental Methods**

*Populations*

ARIC includes data from 2097 African American men and women aged 45-64 years recruited from four US communities: Forsyth County, NC; Jackson, MS; Minneapolis, MN; Washington County; MA. Participants were followed up for up to 5 visits, with DNA methylation derived from visit 2 [1]. The MCCS consists of 5361 men and women aged 40-69 years from the Melbourne region. The study was composed of participants included in six prior nested case-control studies of prostate, colorectal, lung or kidney cancer, urothelial cell carcinoma or mature B-cell neoplasms. Controls were matched on sex, year of birth, country of birth, baseline sample type and smoking status (lung cancer study only) [2]. The LBC consists of individuals born in 1921 and 1936 living in the Lothian region. This study includes the baseline examination of 550 individuals from LBC 1921 (average age 79) and 1091 LBC from 1936 (average age 70) [3]. Lifelines DEEP is a sub-cohort of the LifeLines study consisting of 752 individuals from the Netherlands [4]. The BHS study is a long-term cohort study focused on cardiovascular disease. This study includes 1,485 adult participants from the study (995 non-Hispanic white and 490 African American) recruited during 2006-2010. The RODAM study is a study of Ghanaians recruited from Ghana, London, Amsterdam and Berlin. From the total study population, 736 were included in the EWAS of BMI [5]. The KORA cohort is a population-based health study examining individuals living in the region of Augsburg in Southern Germany. The F3 and F4 surveys were follow up examinations of the original cohort taken in 2004-2005 and 2006-2008, respectively. DNA methylation was measured in 1709 participants from F4 and 285 participants from F3 [6]. The LOLIPOP study is a prospective cohort study of Indian Asian and European men and women recruited in West London, United Kingdom between 2003 and 2008. DNA methylation was measured in 2,680 participants free from type 2 diabetes using peripheral blood collected at enrollment [7]. EPICOR was a nested case-control study from the European Prospective Investigation into Cancer and Nutrition (EPIC)-Italy cohort, recruited during 1994-1998. DNA methylation was measured in peripheral blood collected at enrollment in 292 controls [8].

Data from replication analysis derived from the Women’s Health Initiative. EMPC (n=2200) assessed epigenetic mechanisms underlying associations between ambient particulate matter air pollution and cardiovascular disease within the WHI CT. BAA23 was a case-control study assessing predictors of coronary heart disease (CHD) within the WHI CT (n=1664) and OS (n=443), where cases were identified using eight biomarkers of CHD. AS311 is a matched case-control study of bladder cancer among women within the WHI CT (n = 426) and OS (n = 456). In the WHI, weight was measured on a balance beam scale to the nearest 0.1 kg. Height was measured to the nearest 0.1 cm using a wall-mounted stadiometer.

In replication analyses, relevant covariates included age, race/ethnicity, cell composition, the top three principal components of genetic relatedness, smoking status, clinical trial arm and case-control status (BAA23 and AS311) in the main model and diet quality, physical activity level and socioeconomic status in sensitivity analyses. Age, race/ethnicity (White, AA, Hispanic/Latino, Asian, American Indian, other), and smoking status (current/former or never) were self-reported. Physical activity was self-reported from a self-administered questionnaire. The data were analyzed as total energy expended from light, moderate or vigorous intensity recreational physical activity which includes walking, mild, moderate and strenuous physical activity in kcal/week/kg (MET-hours/week) [9]. Diet quality was estimated using the alternative healthy eating index-2010 (AHEI-2010) [10]. Income status was examined as a proxy for socio-economic status (3 level factor variable reduced from 7 level factor).

**Supplemental Figures**

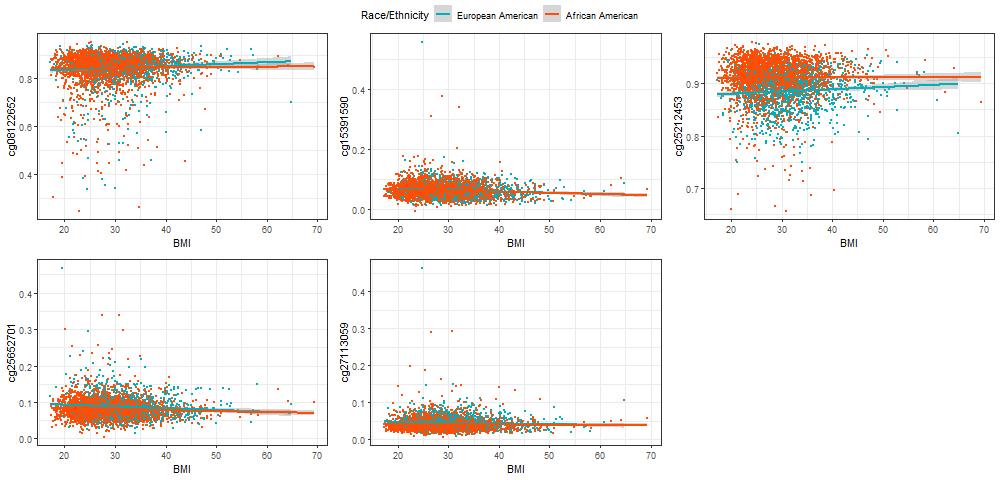

**Figure S1.** Scatter plot of interaction by race/ethnicity in five CpG sites.
